# Epithelial and stromal remodeling in murine Crohn’s disease-like ileitis

**DOI:** 10.1101/2025.11.08.684908

**Authors:** Katherine C Letai, Bianca N Islam, Paola Menghini, Carlo De Salvo, Neha S Khandekar, Bruce W Armstrong, Alka Tomar, Kim Curry, Theresa T Pizarro, Paul J Tesar, Marissa A Scavuzzo, Fabio Cominelli

## Abstract

**Background and Aims:** Inflammatory bowel disease is characterized by progressive epithelial, immune, and stromal dysfunction. Human datasets yield important insights yet are inherently descriptive, requiring animal models for *in vivo* mechanistic interrogation. The SAMP1/YitFc (SAMP) mouse model develops spontaneous Crohn’s disease (CD)-like ileitis while allowing controlled manipulation of pathophysiologic pathways. To understand cell-specific disease-associated transcriptional changes, we defined the single-nucleus transcriptomes of inflamed SAMP ilea using single-nucleus RNA-sequencing (snRNA-seq).

**Methods:** We performed whole-ileum snRNA-seq on SAMP (*n*=4) and control AKR/J (*n*=3) mice using the gut-optimized CitraPrep protocol, generating 58,349 high-quality nuclei across epithelial, immune, and stromal lineages. We corroborated SAMP-associated changes with immunofluorescent staining, Western blotting, spatial transcriptomic, and human single-cell RNA-sequencing data.

**Results:** We observed a primary type 2 immune phenotype in SAMP compared to AKR mice, but most transcriptional changes occurred in stromal and epithelial, not immune, cells. Stromal remodeling included increased fibroblast-derived *Igf1*. In SAMP epithelium, expansion of tuft cells and emergence of SAMP-specific *Pcsk6+* crypt enterocytes were observed and confirmed in human CD patients. Finally, we identified cross-compartmental changes in cell-cell signaling, including enterocyte-to-type 3 innate lymphoid cell (ILC3) communication and enhanced global *Igf1* signaling. These features recapitulate known, as well as novel, characteristics of CD and extend our understanding of disease-associated multicellular networks.

**Conclusion:** This atlas of ileitis-prone SAMP mice provides a high-resolution resource for dissecting conserved and novel mechanisms of inflammation and tissue remodeling. Herein, we uncover disease-associated transcriptional changes conserved in human CD, presenting a translational platform for future mechanistic and therapeutic studies.

## INTRODUCTION

Inflammatory bowel disease (IBD) comprises a group of chronic, relapsing inflammatory disorders of the gastrointestinal tract, with Crohn’s disease (CD) representing one of its more severe and complex forms.[1] While significant strides have been made in understanding IBD pathogenesis, mechanistic insights into cell type-specific alterations remain limited, in part due to the complexity of the human condition and the lack of fully representative animal models.

SAMP1/YitFc (SAMP) mice represent a spontaneous model of CD-like ileitis that share similar features to disease in CD patients, including transmural inflammation, mucosal ulceration, lymphoid aggregates, and occasionally, granuloma formation.[2,3] Importantly, SAMP mice exhibit a polygenic, microbially-influenced disease process without the need for chemical induction or genetic engineering, positioning them as a uniquely valuable model for studying the interplay of genetic susceptibility, epithelial barrier dysfunction, immune dysregulation, and environmental triggers in IBD. We have previously shown that the primary defect conferring ileitis in SAMP mice originates from a nonhematopoietic source.[4] However, the specific cell type and related transcriptomic profile involved in this process have not been determined.

Recent advances in single-cell and single-nucleus RNA sequencing have enabled unprecedented resolution in profiling tissue heterogeneity, allowing researchers to delineate cell-type-specific transcriptional programs and infer intercellular interactions.[5] Single-nucleus RNA sequencing (snRNA-seq) offers practical advantages in tissues that are difficult to dissociate or to preserve intact cells without biasing populations, such as full-thickness intestinal tissues, which is further complicated during disease by fibrosis and inflammatory processes. Large human IBD atlases have relied on separate dissociation steps of isolating epithelial and lamina propria fractions, and when whole tissues were processed, fragile cell types were often lost, leaving mainly fibroblasts and muscle. By contrast, the recently developed CitraPrep protocol preserves RNA integrity across all compartments, enabling high-quality profiling of intestinal tissues that was previously unachievable.[6] This represents a major advance over traditional single-cell workflows, where high concentrations of digestive enzymes often degraded RNA and limited approaches to restricted populations of cells. Despite the growing application of these technologies to human IBD, there remains a paucity of single-cell atlases derived from preclinical models of CD-like ileitis. Generating these atlases is critical for aligning model systems with human disease biology and distinguishing conserved from context-specific pathways.

Despite clear similarities of SAMP mice to CD patients, we have yet to understand how the individual cell types in this model reflect human disease. Data from this atlas aims to enhance our understanding of some of these knowledge gaps. Here, we applied snRNA-seq and spatial transcriptomics to whole ileal tissues from SAMP mice with established disease and compared their gene signature(s) to healthy control AKR mice, generating a multicompartmental transcriptional atlas of chronic ileitis. We defined disease-associated shifts in cellular composition and gene expression of immune, stromal, and epithelial cells. These changes corroborated known key features of SAMP chronic ileitis, including Th2 immune activation and expansion of Paneth cells[7–9], but also revealed novel findings, such as tuft cell expansion and presence of SAMP-specific *Pcsk6+* enterocytes, and highlighted a stromal source of IGF1 in intestinal inflammation.

To understand the translational relevance to these findings, we compared our murine data to a large-scale single-cell RNA-seq dataset from 71 IBD patients and healthy controls[10], revealing shared transcriptional signatures across species. To further dissect how chronic inflammation reshapes tissue organization, we applied interactome analysis across 26 cell types, uncovering ligand–receptor network rewiring between epithelial, immune, and stromal cells. These multicellular circuits highlight how chronic ileitis alters not only cell-intrinsic programs, but also intercellular communication. This work establishes a high-resolution framework for mechanistic exploration and identification of therapeutic targets for chronic ileitis, and supports the use of spontaneous murine models in preclinical IBD research.

## RESULTS

### A global single nucleus transcriptional blueprint of ileitis-prone SAMP ilea unveils disease-associated transcriptional changes in every cell type

To investigate disease-associated transcriptional changes at the single-cell level, we collected whole ileal tissues from four 20-week-old SAMP1/YitFc (SAMP) mice with spontaneous ileitis and three healthy AKR (parental) controls. Gross dissection revealed inflamed, thickened ileal tissue in SAMP mice (online supplemental figure 1A). Histologic scoring confirmed severe ileal inflammation in SAMP compared to AKR mice (figure 1A). We isolated nuclei using the recently developed gut-optimized CitraPrep method[6] (online supplemental figure 1B), allowing snRNA-seq of these samples (figure 1B). Filtering out low quality nuclei and doublets produced 58,349 high-quality single-nuclear transcriptomes (figure 1C–E, online supplemental figure 1C-E). Based on unsupervised clustering and marker expression, nuclei were divided into three major tissue compartments: epithelial (81%), immune (10%), and stromal (9%) (figure 1F–G, online supplemental figure 1F, online supplemental table 1), exhibiting well-established cell-specific gene markers (*e.g*., *Vil1, Ptprc, Acta2*, respectively) (figure 1H). This compartmental distribution in mouse ileal tissues approximates the cellular composition reported in human single-cell datasets of healthy ileum, wherein ∼80–90% of cells are epithelial, 8– 15% immune, and 5–10% stromal, supporting the biological relevance of our murine dataset.[11–13] To ensure statistical robustness, we applied pseudobulk differential expression analysis that aggregates nuclei by biological replicate, a more stringent approach than single-cell–level testing alone. This analysis revealed that in almost every cell type, more genes were downregulated than upregulated in SAMP compared with AKR mice, similar to patterns seen in human ileitis (figure 1I).[10]

**Figure 1:**
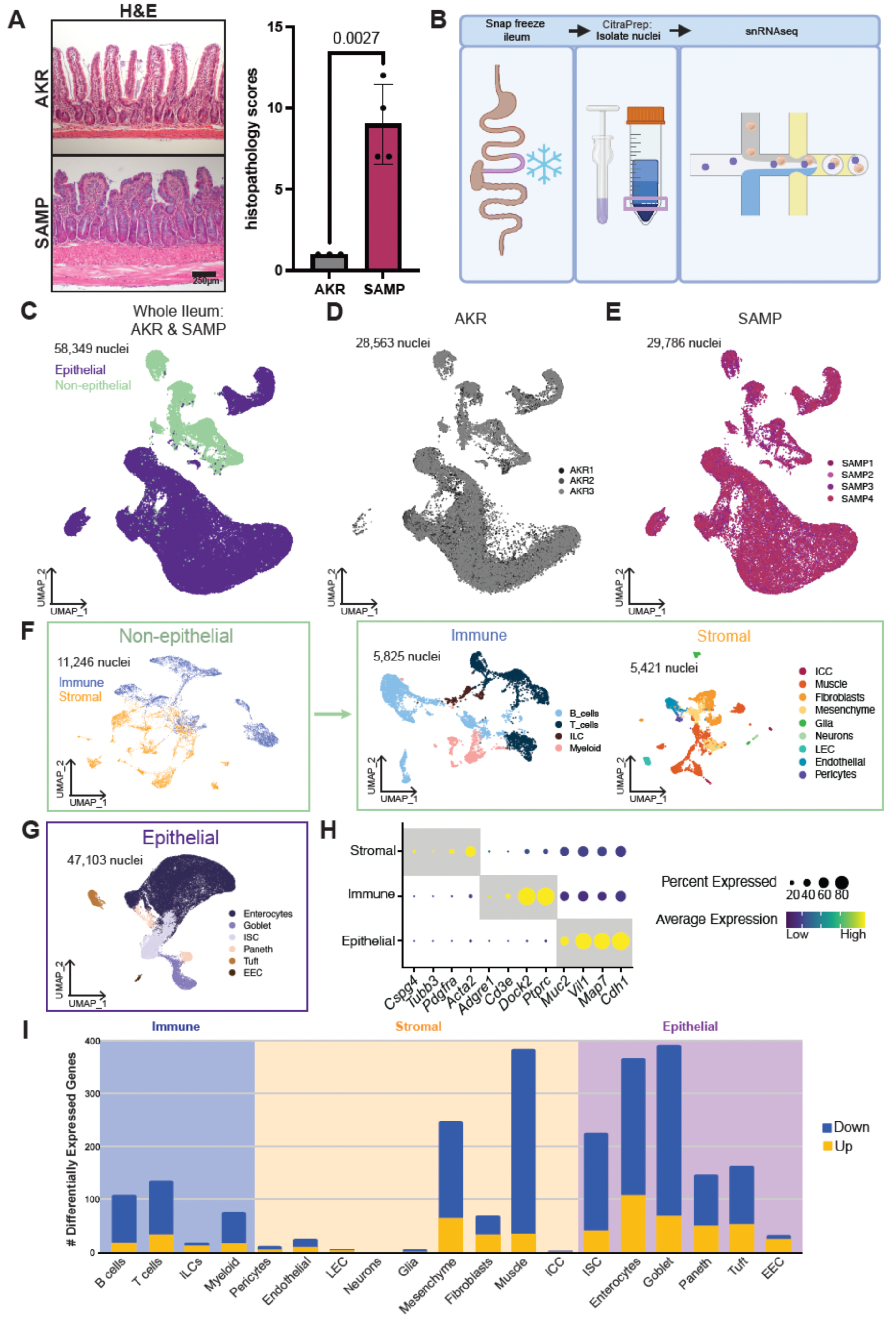
Single nucleus transcriptional blueprint of the murine ileum across epithelial, immune, and stromal compartments. (A) Left, representative H&E images illustrate hallmark features of ileitis in SAMP mice, including villous distortion, crypt architectural disruption, and prominent leukocytic infiltration in the lamina propria and submucosa. Right, blinded histopathological scoring of ileal inflammation in SAMP versus AKR (*n*=3-4 per group; exact P shown on plot). (B) Schematic of nuclei isolation and sequencing process. Ileal tissues (10 cm) were harvested from control AKR (*n*=3) and SAMP (*n*=4) mice, and nuclei isolated using the CitraPrep protocol for downstream snRNA-seq. (C–E) UMAPs of whole-atlas nuclei from all mice (C), AKR only (D), and SAMP only (E). Nuclei are colored by cellular compartment in (C) and by biological replicate in (D–E), and visualized with Uniform Manifold Approximation and Projection (UMAP) (*n*=7 mice, 58,349 nuclei after QC). (F) Non-epithelial cells were split into immune and stromal sub-atlases, with UMAPs colored by cell type. (G) Epithelial nuclei are colored by epithelial cell type and visualized by UMAP. (H) Expression of compartment-specific markers. Dot plot shows canonical markers (rows) across compartments (columns); dot size indicates the fraction of nuclei expressing the gene and color reflects average scaled expression. (I) Bar graph reporting the number of up- and down-regulated genes in SAMP versus AKR across major epithelial, immune, and stromal lineages (cell type groupings as shown), using thresholds defined in Methods.

### Dominant type 2 immune activation in SAMP

Next, we sought to identify differences in the immune environment between these two strains at the single cell level. We resolved the immune sub-atlas by strain and identified the major lymphoid and myeloid lineages in both AKR and SAMP mice, recovering 3,077 immune nuclei from AKR and 2,748 from SAMP (figure 2A-B). We defined specific cell subtype identities by unsupervised clustering and marker expression, including B and plasma cells, CD4+ and CD8+ T cells, ILCs, dendritic cells, macrophages, eosinophils, mast cells, and cytotoxic lineages (figure 2A-B; online supplemental figure 2A). We further confirmed cell type assignments by visualizing expression of known markers for each identity (online supplemental figure 2B). Functional enrichment analysis of upregulated genes comparing all immune cells in SAMP with those of AKR suggests generalized immune activation and cytokine signaling in SAMP, encompassing a variety of inflammatory response types (figure 2C, online supplemental table 2). Furthermore, the CD-associated proinflammatory cytokines, *Il1a, Il1b, Tnf*, and *Il23a*, exhibited altered expression patterns across immune cell subpopulations in SAMP versus AKR mice (figure 2D, online supplemental figure 2C), confirming earlier studies.[2] A feature plot of *Ccl21a* and *Ccl19* highlights a statistically significant (adj p=0.00059), and a trend towards, decreased expression, respectively, in ilea from SAMP, both of which have been previously shown to be downregulated in SAMP mice (online supplemental figure 2D)[14]. Similar to human studies, the immune compartment exhibited fewer transcriptional changes than stromal and epithelial cells (figure 1I).[10] We were therefore particularly interested in assessing changes in cell type proportion between SAMP and AKR immune cells (figure 2E). Type 2 immunity orchestrators ILC2s (q=7.3e-4), along with eosinophils and mast cells (q=1.2e-2), were significantly expanded in SAMP, confirming prior studies establishing SAMP mice as a predominant Th2-driven model.[7,15,8] Strain-stratified feature overlays further indicated a shift toward type 2 polarization in SAMP, reflected by higher *Gata3* relative to *Tbx21* signals and an increased proportion of *Gata3+Tbx21-* versus *Tbx21+Gata3-* cells (Fig. 2F). Concordantly, the Th2 cytokines, *Il5, Il4,* and *Il13*, exhibited higher expression in SAMP immune cells than AKR (online supplemental figure 2C). These analyses defined the immune landscape of the SAMP ileum and detailed a Th2-leaning inflammatory milieu, substantiating previous characterization of the SAMP mouse and opening the door for subsequent epithelial–immune–stromal interaction studies.

**Figure 2:**
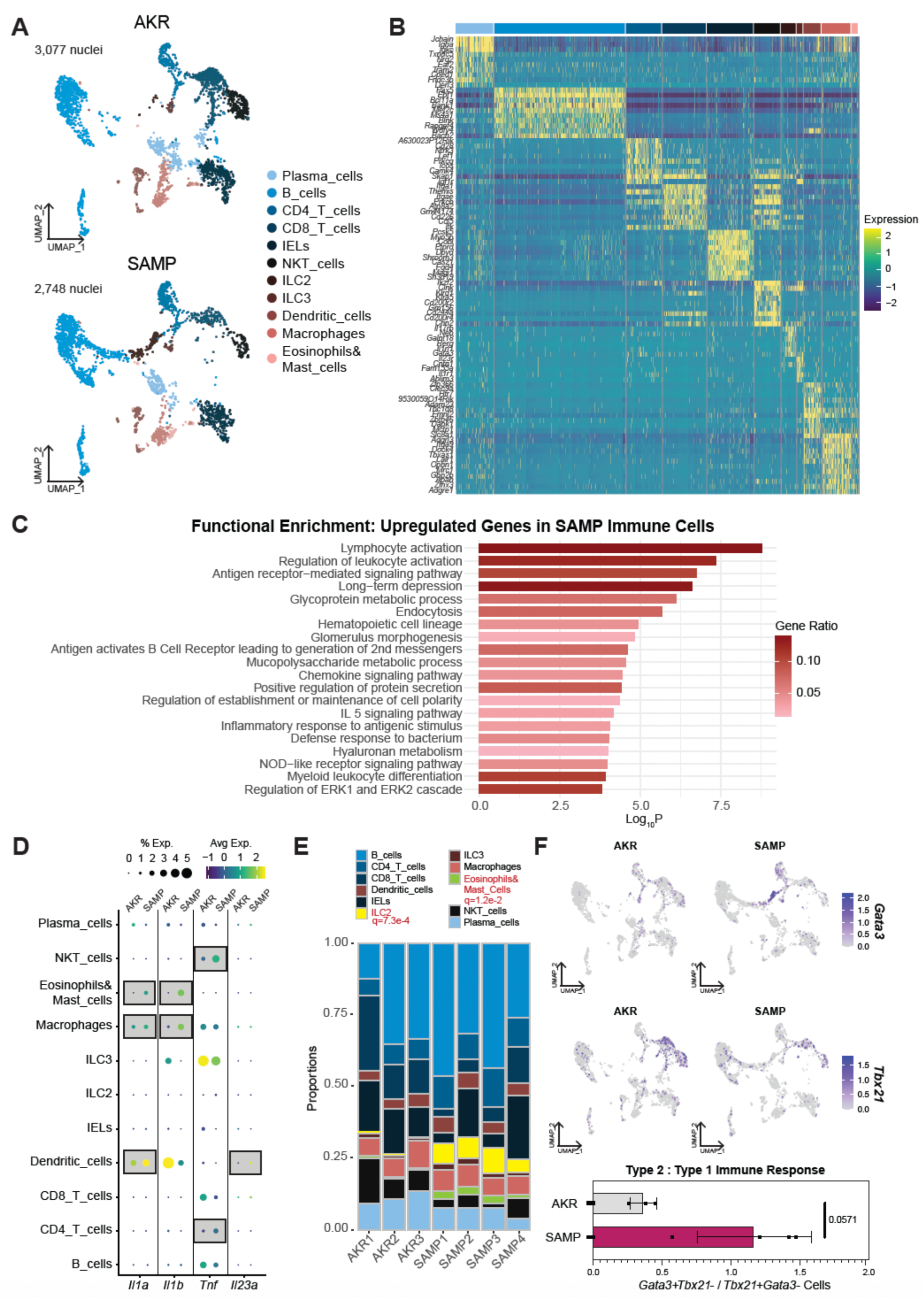
Type 2 immune activation in SAMP ileum. (A) Immune atlas UMAPs stratified by strain, showing AKR (control) and SAMP (disease) nuclei colored by cell type. Unsupervised clusters shown in online supplemental figure 2A. (B) Cluster-defining expression heatmap for immune lineages, displaying top 10 transcripts distinguishing each population in rows, and each column representing an individual nucleus, with cell types clustered together and labeled by color above. Full marker lists are provided in online supplemental table 1. (C) Functional enrichment of genes upregulated in SAMP immune cells, showing Gene Ratio and –log10P as shown for each represented functional term. (D) Dot plot showing *Il1a, Il1b, Tnf,* and *Il23a* across immune cell types, stratified by strain. Dot size indicates percent of cells expressing the transcript, and color indicates relative scaled expression. (E) Per-sample immune cell-type composition (AKR1–3; SAMP1–4), shown as a proportion of the full immune cell population. q-values <0.05 are displayed, with labels for expanded populations in red and diminished populations in blue. (F) *Gata3* and *Tbx21* overlaid on strain-specific immune UMAPs, with color representing normalized expression compared with all genes across all immune cells. Below, quantification of type 2–biased (*Gata3+ Tbx21-*) versus type 1–biased (*Tbx21+ Gata3-*) cells (Type 2:Type 1 ratio shown on plot).

### SAMP fibroblasts exhibit profibrotic and proinflammatory transcriptional activation

Beyond immune changes, we also asked whether the stromal compartment differed in gene expression patterns between SAMP and AKR mice. Consolidation of unsupervised clusters of stromal cells based on similar transcriptomic profiles indicated representation of mesenchymal, endothelial, and neural lineages in both AKR and SAMP ilea (figure 3A-B, online supplemental figure 3A-B). Given the known disease-related changes in fibroblasts in IBD, we asked whether SAMP fibroblasts transcriptionally shift towards an activated subtype.[16,17] As expected, fibroblasts from SAMP mice displayed significant upregulation of a cohort of transcripts (*Fn1*, *Postn*, *Timp1*, *Ccl2*, *Pdgfra*, *Lox*, *Tnc*, *Wnt2b*, *Il11*, and *Cxcl1*) related to inflammation and fibrosis compared to AKR controls (p = 9.7e–15, figure 3C), confirming a robust fibrotic phenotype in SAMP ilea[18,19] A key upregulated gene revealed by differential expression analysis was *Igf1* (figure 3D-E, online supplemental table 2). In Crohn’s patients, *IGF1* has been implicated in intestinal extracellular matrix (ECM) collagen regulation, aggravation of fibrosis, and facilitation of malignant transformation.[20,21] To understand whether fibroblasts may be a source of IGF1 protein, we performed multiplexed immunofluorescent staining and confirmed elevated IGF1 in COL1A1+ACTA2+ fibroblasts in SAMP ilea compared with AKR (figure 3F). *Igf1* upregulation is just one of many changes in fibroblast gene expression, so we next asked what other functional changes are implied by these changes. Functional enrichment analysis revealed upregulation of pathways involved in extracellular matrix remodeling, actin filament-based movement, and cell migration, while downregulated programs included interferon signaling and proteoglycan metabolism (figure 3G). Given that muscle cells exhibit the greatest amount of transcriptional change out of stromal cells (figure 1I), as well as the dramatic thickening of the muscle layer observed on histology (figure 1A), we also wanted to understand which transcripts were changing in myocytes of SAMP mice. Notably, we found downregulation of *Calm1* and *Calm3* in SAMP muscle cells, which both code for Calmodulin, a critical protein for proper muscle contraction (online supplemental figure 3C, online supplemental table 2). Functional enrichment analysis (online supplemental figure 3D) indicates mostly metabolic and inflammatory changes, suggestive of adaptive myocytes adapting to an inflammatory environment. Aside from mesenchymal cells, all other stromal cell types, including enteric nervous system cells, exhibited minimal transcriptional differences between SAMP and AKR strains (figure 1A).

**Figure 3:**
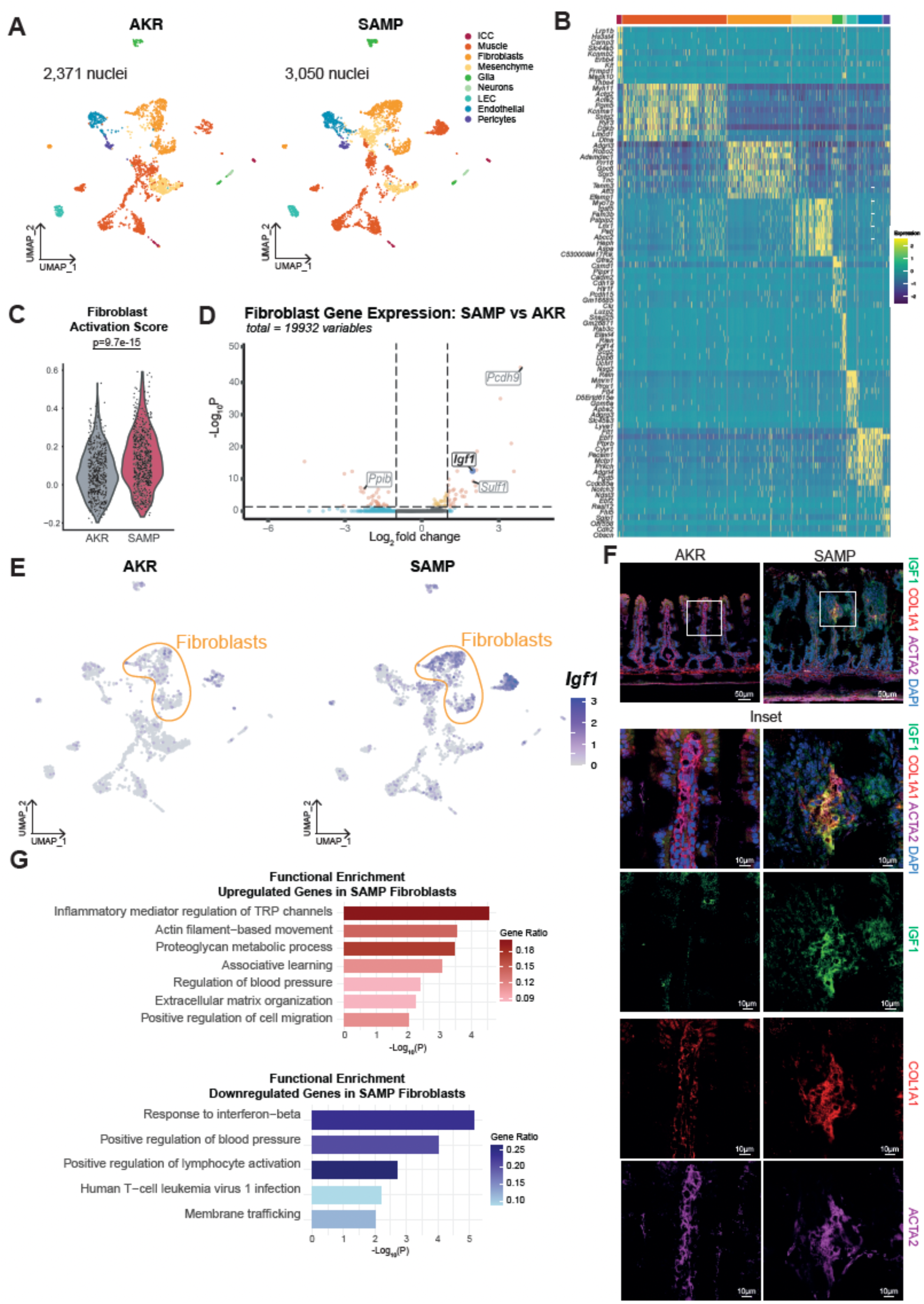
Stromal remodeling in SAMP ilea indicates fibroblast activation and increased IGF1 expression. (A) Stromal nuclei from AKR and SAMP animals plotted separately by UMAP and colored by cell type to visualize population differences. Unsupervised clusters shown in online supplemental figure 3A. (B) Top 10 transcripts enriched in each stromal cell type. Individual transcripts are shown in rows, cell types are clustered together in columns with color annotation above. Full marker lists are provided in online supplemental table 1. (C) Fibroblast Activation Score in AKR and SAMP, p=9.7e-15. Module score was calculated using average expression of the *Fn1*, *Postn*, *Timp1*, *Ccl2*, *Pdgfra*, *Lox*, *Tnc*, *Wnt2b*, *Il11*, and *Cxcl1*. (D) Differential gene expression in SAMP fibroblasts compared with AKR visualized as a volcano plot. Log_2_ fold change (x-axis) and -Log_10_P calculated using DESeq2 pseudobulk analysis. (E) *Igf1* expression in AKR and SAMP stroma plotted separately for visualization in UMAP space. Color represents normalized expression compared with all genes across all stromal cells. Fibroblasts are circled. (F) Immunofluorescent staining of IGF1 in ACTA2+COL1A1+ fibroblasts shows an increase in SAMP compared with AKR ileum. (G) Functional enrichment analysis of upregulated and downregulated genes in SAMP fibroblasts. –log10P for each represented functional term is plotted, colored by gene ratio.

### Expansion of secretory lineages and emergence of *Pcsk6+* enterocytes in ileitis

We next asked what transcriptional changes occur in the diseased epithelium. Unsupervised clustering of 47,103 epithelial nuclei identified six canonical cell types: enterocytes, goblet cells, intestinal stem cells (ISCs), Paneth cells, tuft cells, and enteroendocrine cells (EECs) (figure 4A-B; online supplemental figure 4A-B). In our analyses, the proportion of SAMP epithelial cells was biased towards the secretory lineage, (q=4.6e-5, online supplemental figure 4C). Specifically, SAMP mice exhibited a significant expansion of Paneth cells, consistent with prior reports[22–24] and newly reported expansion of tuft cells relative to AKR controls (figure 4C), with tuft cell enrichment validated by DCLK1 immunostaining (p=0.0083, figure 4D-E).

**Figure 4:**
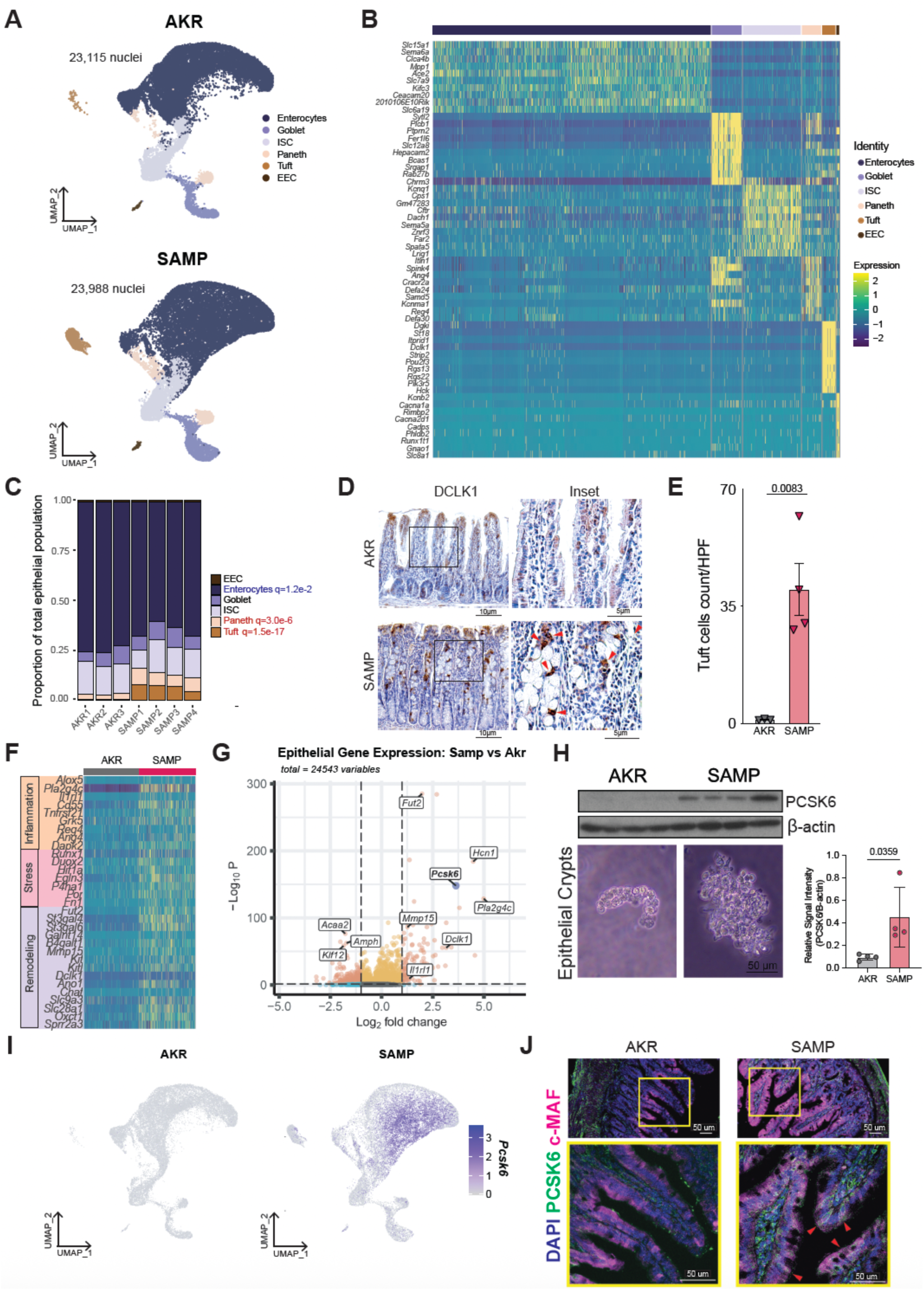
Secretory expansion and PCSK6+ enterocyte in the SAMP epithelial compartment. (A) Epithelial nuclei from AKR and SAMP animals plotted separately and colored by cell type to visualize population differences. Visualized by UMAP. Unsupervised clusters shown in online supplemental figure 4A,D. (B) Top 10 transcripts enriched in each epithelial cell type. Individual transcripts are shown in rows, cell types are clustered together in columns with color annotation above. Datasets are provided in online supplemental table 1. (C) Fraction of nuclei of each cell type out of all epithelial nuclei per sample. All q-values <0.05 listed in red for populations expanded in SAMP, and blue for populations decreased. (D) Immunohistochemistry of the tuft cell marker DCLK1 in both AKR and SAMP tissues, with inset showing example fields used for quantification. Red arrows mark nucleated tuft cells. (E) Quantification of tuft cell numbers normalized per high power field (20X) in AKR and SAMP ilea. Each point represents one animal, with error bars representing standard error and p value (0.083) calculated from a two-sample t-test. (F) SAMP tissues exhibit disease-related transcriptional changes. Heatmap of genes involved in inflammation, stress, and tissue remodeling. Individual transcripts are shown in rows, organized by category, and cells are clustered by disease status in columns with color annotation above. (G) Differential gene expression in SAMP epithelial cells compared with AKR visualized as a volcano plot. Log_2_ fold change (x-axis) and -Log_10_P calculated using DESeq2 pseudobulk analysis to retain biological replicate. Select genes of interest annotated, with further investigation focused on *Pcsk6* (bold, purple data point). (H) On the left, Western blot analysis of PCSK6 in isolated ileal crypt-villus units from AKR and SAMP, with representative brightfield images from crypt isolation shown below. Each column represents protein from a different mouse, with ß-actin shown in the second row as a loading control. On the right, quantification of PCSK6 in each mouse normalized by ß-actin. (I) *Pcsk6* expression in AKR and SAMP epithelium plotted separately for visualization of this differentially expressed gene (DEG) in UMAP space. Color represents normalized expression compared with all genes across all epithelial cells. (J) Immunofluorescent staining of PCSK6 in epithelial crypt-villus from AKR and SAMP ileum, images taken at 20X are shown at the top, with high power images (60X) in the yellow boxed inset for better visualization. Red arrows mark example PCSK6+ enterocytes.

To identify the most salient transcriptional changes in the SAMP epithelium, we performed differential expression analysis comparing all epithelial cells across strains (figure 4F-G, online supplemental table 2). Functional enrichment analysis highlighted that SAMP-derived epithelial cells exhibited upregulated markers associated with epithelial inflammation, stress, and remodeling, including *Reg4*, *Ang4*, *Cd55*, and *Duox2* (figure 4F). One of the top upregulated genes in SAMP was *Pcsk6*, encoding proprotein convertase subtilisin/kexin type 6 (figure 4G). PCSK6 is a calcium-dependent serine protease that cleaves proteins to their active forms and was previously shown to promote Th1 differentiation in colitis by activating STAT1.[25] Western blot of epithelial crypt-villus units[26] confirms the upregulation observed in snRNA-seq is maintained at the protein level (Fig. 4H), with the potential to have functional effects. We next visualized *Pcsk6* across all epithelial cell subtypes, revealing that expression is primarily restricted to a unique subset of enterocytes only present in SAMP tissues, Enterocyte7 (figure 4I, online supplemental figure 4D-E). Functional enrichment analysis of the top 100 marker genes of this enterocyte cluster suggested changes to metabolic and immune-related programs (online supplemental figure 4F), including multiple terms related to extracellular matrix production and glycosylation. Immunofluorescent staining of SAMP and AKR tissues confirmed the appearance of PCSK6 protein in a subset of SAMP enterocytes (c-MAF+) (figure 4J), illustrating the specificity of this enterocyte subtype in the diseased ileum. Beyond *Pcsk6,* SAMP enterocytes also upregulated *Il1rl1* (ST2) at both the transcript and protein levels, suggesting IL-33 responsiveness and a further contribution to epithelial–immune signaling (online supplemental figure 4G-H).

### Spatial transcriptomics and human single cell validation of the SAMP ileal atlas

To contextualize the potential functional changes of SAMP epithelial cells, we wanted to know whether epithelial programs identified by snRNA-seq were spatially localized in mouse ilea and conserved in human IBD. Due to the discontinuous, patchy nature of disease in SAMP mice, whole ilea harvested for snRNAseq contained nuclei from both inflamed (cobblestone-like)[27] and noninflamed regions that cannot be distinguished from each other using this approach. GeoMx enables us to assess compartment-level transcriptomes in AKR and SAMP mice, segregating SAMP tissue into cobblestone (C)/inflamed versus non-cobblestone (NC)/noninflamed regions (figure 5A, online supplemental figure 5A). Spatial analysis of the epithelial compartment revealed higher transcript counts in SAMP mice for *Pcsk6, Dclk1, Il1rl1, Pla2g4c, Fut2,* and *Mmp15* (figure 5B), mirroring the snRNA-seq data (figure 4G). Cobblestone and non-cobblestone regions exhibited similar trends in expression profiles when compared to AKR, suggesting many of the most relevant transcriptional changes in SAMP ilea occur even in the absence of inflammation. Atlas-derived DEGs were also differentially expressed in the immune and stromal compartments of SAMP and AKR mice, with cobblestone and non-cobblestone regions again showing similar expression patterns (online supplemental figure 5B, online supplement table 1). Together these spatial transcriptomic data not only confirm our snRNAseq findings, but also indicate that many disease-associated transcriptional changes occur globally in the SAMP ileum.

**Figure 5.**
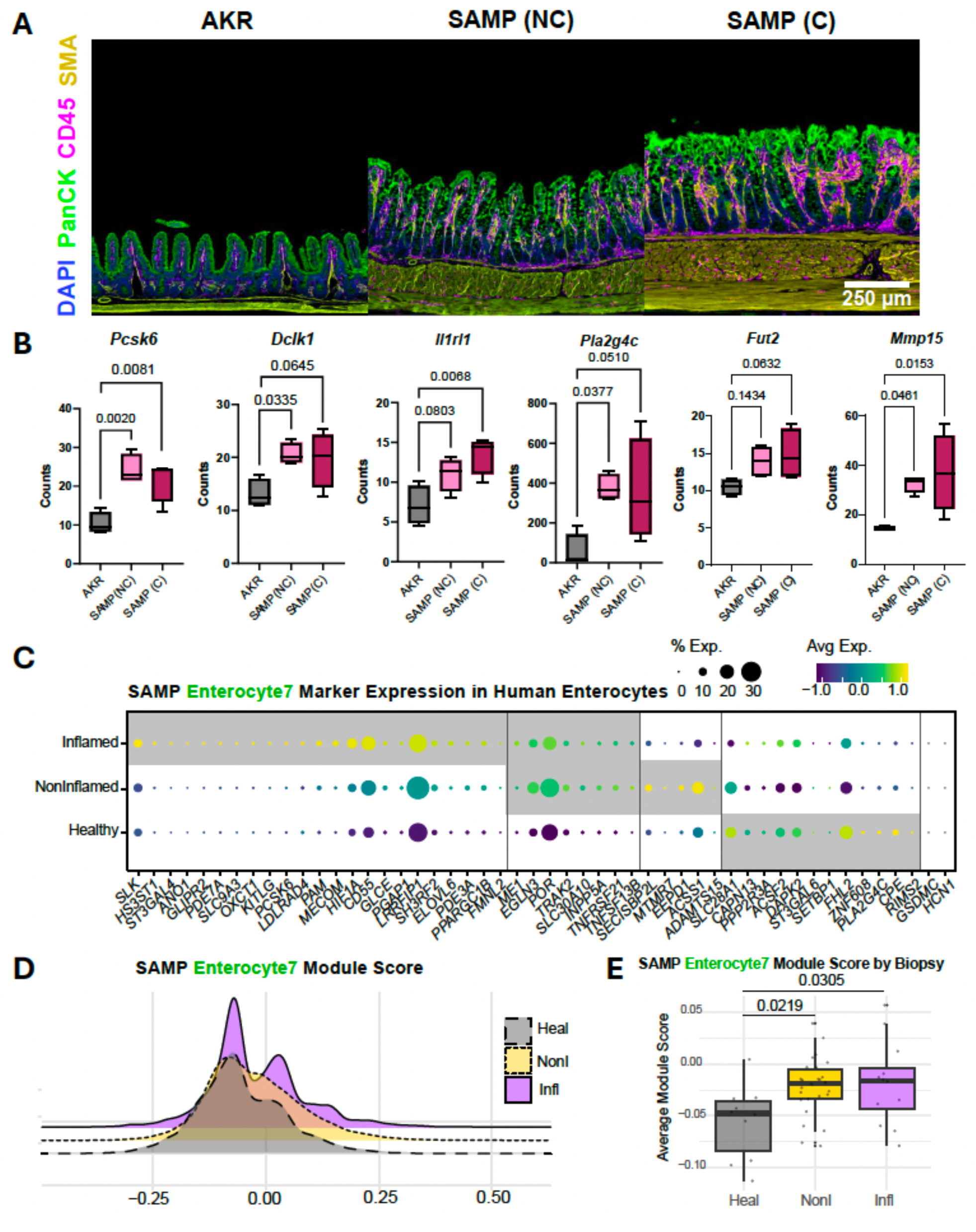
Spatial transcriptomics and human single-cell data validate epithelial programs from the SAMP ileal atlas. (A) Mouse spatial transcriptomics of ileum comparing AKR and SAMP regions of interests. Representative images showing tissue architecture in AKR and SAMP (both non-cobblestone and cobblestone regions with nuclei (DAPI), epithelium (PanCK), immune infiltrates (CD45), and stromal compartment (SMA) overlays, scale bar 250 um. (B) Box plots of spatial expression for PanCK+ genes across AKR, SAMP NC (non-cobblestone), and SAMP C (cobblestone) regions; exact *P*-values displayed. (C) Dot plot of SAMP Enterocyte7 marker genes in human ileal enterocytes across healthy, noninflamed IBD, and inflamed IBD cohorts; dot size denotes percent expressing and color indicates average scaled expression. Grey boxes highlight (left-to-right) transcripts elevated primarily in Inflamed tissue, similarly elevated in Inflamed and Noninflamed, elevated in Noninflamed only, and similar or elevated in healthy cells. (D) Ridge plot of Enterocyte7 module score distribution across human enterocytes for healthy, noninflamed, and inflamed groups. (E) Per-sample Enterocyte7 module scores in the human dataset, displaying individual patients with group medians/interquartile ranges; exact P values reported on panel.

We next asked whether the enterocyte changes we observed in the SAMP model were also present in patients with CD. We assessed expression of the top 50 murine Enterocyte7 gene markers in a single cell RNAseq dataset[10] derived from human ileal-specific CD. At the gene level, 72% of Enterocyte7 markers displayed increased expression in enterocytes of noninflamed or inflamed IBD relative to healthy controls (figure 5C). We next calculated module scores of the Enterocyte7 marker panel in human enterocytes. Enterocyte7 score was elevated in noninflamed and inflamed human enterocytes at both the single cell (figure 5D) and at the biopsy levels (figure 5E) compared with scores derived from healthy ileum, in line with our finding that alterations in SAMP-distinguishing gene expression occur in both affected and unaffected epithelium. This congruency again supports the translational relevance of the Enterocyte7/*Pcsk6+* enterocyte signature in CD-specific ileitis.

### Cross-compartment interactome analysis reveals modified cell–cell communication in chronic ileitis

To assess whether transcriptional remodeling alters intercellular communication, we performed ligand–receptor interactome analysis across 26 cell types.[28] Based on expression of ligands and receptors, this analysis predicted overall interaction strength between cell types. One of the cell-cell communication relationships predicted to be strongest in SAMP compared with AKR ileum is enterocyte signaling to ILC3s (figure 6A), confirming the importance of this ILC subset in the pathogenesis of SAMP ileitis.[29] Both disease-only and disease-upregulated interactome pairs were identified between enterocytes and ILC3s (figure 6B). Interestingly, examination of ligand expression across enterocyte subclusters revealed that the ILC3-activating signal, *Kitl*, is expressed specifically in the *Pcsk6*+ Enterocyte7 cluster (figure 6C), providing additional evidence that this SAMP-restricted enterocyte state may promote tissue inflammation.

**Figure 6:**
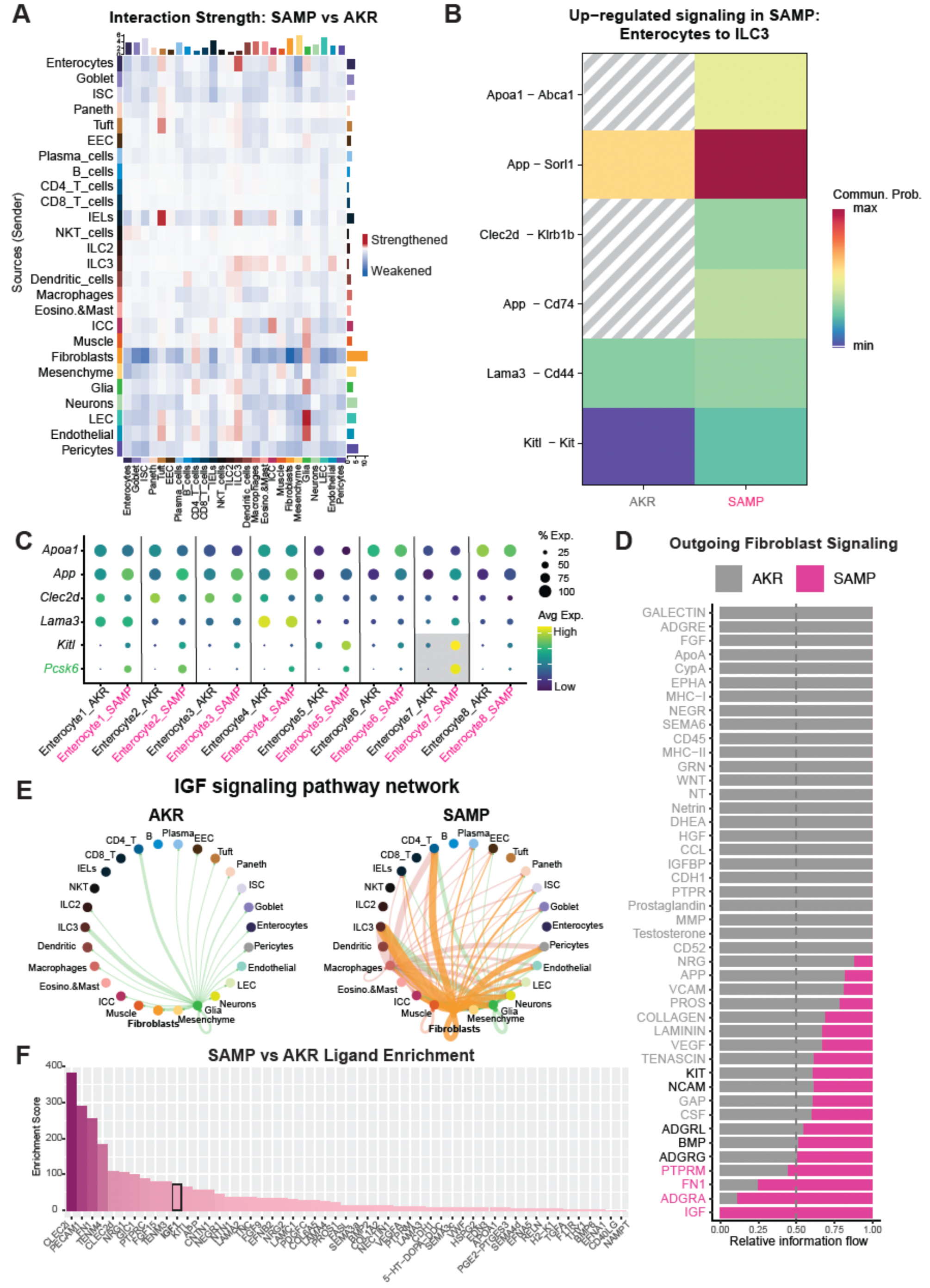
Cross-compartment interactions in SAMP ileum reveal tissue-wide rewiring of inflammation and repair programs. (A) Heatmap showing the strength of interactions between all cell types in SAMP vs AKR animals, predicted by CellChat. Source cells plotted on the y-axis, and receiving cells are on the x-axis. Height of bars represents the sum of the absolute values of the heatmap values in that row (righthand bars) or column (top bars) (higher bar means more change for that cell type). (B) Communication probabilities for changed interactions from enterocytes to ILC3s. Probabilities in AKR are listed on the left, and SAMP on the right. Color represents probability, with red as high and blue low. All interaction probabilities have p-values <0.01 when compared between SAMP vs AKR. Grey lines indicate no signaling predicted for that ligand-receptor pair and strain. (C) DotPlot showing expression of the predicted ILC3-sensed ligands transcribed by enterocytes (rows), in SAMP vs AKR cells across all 8 enterocyte subtypes in our dataset (columns). *Kitl* exhibits the most dramatic increase in SAMP vs AKR, particularly in Enterocyte7, alongside *Pcsk6* (green). (D) Relative information flow of the fibroblast outgoing signaling network. Information flow is defined as the sum of communication probability among all pairs of cell groups, plotted here by condition as a proportion of the total communication across both conditions (AKR+SAMP). Pathways in grey are enriched in AKR, and those in pink are enriched in SAMP. (E) Circos plots showing IGF signaling in AKR vs SAMP. Lines are colored like the cell type sending the signal, connecting to cell types predicted to receive the signal, with outgoing fibroblast signals highlighted in purple. (F) Enrichment score for all ligands enriched in SAMP (across all cell types), including IGF (black box).

To predict which cell types exhibited the most change in cell-cell signaling in SAMP tissues compared with AKR, the tissue-wide interactome analysis summed absolute values in the changes of interaction strengths across all ligand-receptor pairs (figure 6A). Fibroblasts had the greatest amount of predicted change in outgoing signaling, mostly weakening their outgoing interactions in SAMP compared to AKR ileum (figure 6A,D). Examining the strength of predicted fibroblast-derived signaling pathways active in SAMP versus AKR tissues revealed IGF was the only pathway predicted to be active exclusively in SAMP fibroblasts (figure 6D). Visualizing IGF signaling across all cells in our dataset demonstrated that IGF signals were initiated only by glia in AKR mice (figure 6E). In contrast, there was extensive rewiring of IGF signaling predicted in SAMP, with fibroblasts as the most prominent source, as well as ILC3s and CD4+ T cells (figure 6E). The interactome predictions indicated that the IGF signal was received by almost every cell type and further suggested IGF was one of the most enriched signaling pathways in SAMP across all cell types (figure 6E). Since we observed putative rewiring of IGF in the SAMP tissue across many cell types, we next set out to determine broad ligand changes in the inflamed gut. Tissue-wide analysis of ligand transcription predicted SAMP enrichment of not only IGF1 secretion, but also CLEC2i, PECAM1, and others, emphasizing that enterocyte-to-ILC3 and IGF signaling are only two of many likely cell-cell signaling changes in SAMP compared to AKR (figure 6F). These putative cross-compartmental interactions indicate that epithelial, immune, and stromal cells all actively modulate their microenvironment during chronic disease. This highlights the need for robust experimental animal models to dissect the cellular and molecular choreography in IBD pathogenesis.

## DISCUSSION

Using snRNA-seq of whole ileal tissues from SAMP mice with spontaneous chronic ileitis, we delineated compartment-wide transcriptional remodeling spanning epithelial, immune, and stromal cell lineages. Our data provide molecular, cell-specific evidence for prior observations of the SAMP pathology, extend the understanding of disease-associated cell states, and uncover novel transcriptional programs that may be relevant to the pathogenesis of CD.

Although CD is classically regarded as an immune-mediated disorder, our results reveal that the most profound transcriptional changes in the SAMP model occur in the epithelial and stromal compartments. This was initially unexpected, given the known importance of T cell–driven inflammation in this model.[2,7] However, this finding aligns with prior work demonstrating that early epithelial alterations occur in SAMP mice, prior to the onset of ileitis, and that epithelial-derived IL-33 promotes early ILC2 expansion in a NOD2- and microbiota-dependent manner during the development of ileitis.[4,8] Our current data extend these findings by showing the presence of specific enterocyte subtypes in SAMP mice. In addition, we show upregulation of ST2+ enterocytes and a larger ILC2 population compared to controls. Fibroblasts from SAMP mice exhibited robust activation signatures, including upregulation of *Igf1*, consistent with a phenotype that promotes both fibrosis and epithelial repair. These findings were validated at the protein level by multiplex immunofluorescence. Gene enrichment analysis suggested that fibroblasts in chronic ileitis adopt a pro-repair and migratory state, activating pathways implicated in ECM reorganization and growth factor signaling. These stromal dynamics provide a unique opportunity to interrogate fibroblast subsets as potential therapeutic targets in chronic intestinal inflammation and IBD-associated fibrosis.

In this study, we observed significant expansion of both Paneth and tuft cells in the SAMP ileum. While increased Paneth cells are consistent with previous reports in this model,^22^ tuft cell expansion diverges from human IBD, where tuft cells are typically depleted in inflamed tissue.^10,29–31^ This distinction may reflect species-specific responses to chronic inflammation or distinct temporal phases of epithelial remodeling. Humans are rarely biopsied until they present with symptoms, raising the possibility that tuft cell expansion represents an early epithelial response captured in this experimental model, but missed in human disease. Both temporal and species-specific explanations, therefore, remain plausible. In parallel, we identified a transcriptionally distinct population of *Pcsk6+* enterocytes, enriched in the villus-crypt regions of SAMP mice. These cells expressed genes involved in secretory stress, antimicrobial defense, and wound healing, suggesting an inflamed crypt-adapted state. Importantly, recent work has shown that PCSK6 promotes colitis progression by binding to and activating STAT1, thereby enhancing Th1 differentiation and M1 macrophage polarization, while compromising tight junction integrity and barrier function *in vivo.*[25] In murine models of DSS-induced colitis, PCSK6 deficiency attenuated inflammation, improved epithelial barrier integrity, and reduced proinflammatory immune polarization. The increased expression of *Pcsk6* in inflamed villus-crypts in our model indicates a broader role for epithelial-derived PCSK6 in shaping mucosal immunity and amplifying local inflammation. Together, these data position PCSK6 as an important candidate mediator of epithelial–immune crosstalk in chronic ileitis, and support further functional interrogation *in vivo*.

Spatial transcriptomic mapping confirmed that increased *Pcsk6* was present in both inflamed and noninflamed SAMP tissue, suggesting a transcriptional change that may precede lesion formation. Notably, parallel analysis of human ileal CD revealed conserved upregulation of the murine Enterocyte7 marker panel, underscoring their translational relevance and establishing SAMP as a preclinical platform for the discovery of human-relevant novel therapeutic targets.

While this study provides a comprehensive snRNA-seq atlas of chronic ileitis in the SAMP model, several technical considerations warrant discussion. First, the use of snRNA-seq, while optimized for frozen tissue, has limitations in capturing predominantly cytoplasmic transcripts. This may be a strength when assessing context-specific changes, in that nuclei sequencing captures nascent transcripts, representing active transcription, rather than long-lived cytoplasmic mRNAs. However, inflammatory mediators, such as cytokines are often underrepresented in nuclear transcriptomes, which may obscure certain facets of immune activation.[33] However, key inflammatory programs were supported by gene module scoring, pathway enrichment, and independent protein validation. Second, we observed limited representation of granulocytes, a known limitation of snRNA-seq, due to their particularly fragile nuclei. As such, granulocyte dynamics are likely underestimated and should be further explored using complementary approaches. Despite these caveats, our dataset includes 58,349 high-quality nuclei from whole ileum tissue across seven animals (*n*=4 SAMP, *n*=3 AKR), yielding high statistical power for differential gene expression, cell-type classification, and cross-compartment interactome analyses.

The identification of disease-associated transcriptional programs across epithelial, immune, and stromal compartments in the SAMP model offers a foundation for translational research aimed at disrupting chronic intestinal inflammation. Our interactome analysis uncovered expanded ligand–receptor signaling from epithelial and fibroblast compartments involving growth factors (*Igf1, Vegfa, Bmp2*), laminins (*Lama3, Lama4*), and adhesion molecules (*Kitl, App*), many of which are implicated in tissue remodeling, angiogenesis, and immune cell recruitment. Several of these pathways are already targeted in cancer or fibrosis therapeutics and may represent underexplored opportunities in IBD.[34–37] For example, fibroblast-derived IGF1 has been shown to regulate epithelial repair and stromal-immune crosstalk[38], and our protein validation confirms its spatial localization to fibroblasts during ileitis. The widespread induction of IGF1 signaling across many cell types in the SAMP ileum also may suggest a compelling therapeutic target, with the ability to impact not just one, but several populations in the chronic ileitis microenvironment.

These insights suggest that the SAMP single-nucleus atlas can serve not only as a mechanistic resource but also as a preclinical platform to nominate and functionally test therapeutic candidates that operate across mucosal compartments. Interventions that target both effector T cells and epithelial/stromal drivers may be especially effective in disrupting the self-sustaining loop of inflammation and maladaptive repair characteristic of CD. Spatial and cross-species validation supports these pathways as conserved drivers of mucosal injury and repair, rather than model-specific features. Integrating spatial and human IBD data into the SAMP framework enhances its translational value and prioritizes clinically relevant targets.

In summary, this study defines the multicellular architecture of chronic ileitis in SAMP mice, reveals novel disease-associated transcriptional programs, and lays the groundwork for targeted investigations of mucosal inflammation and repair. The inclusion of all major compartments in a single atlas enhances its utility as a reference for future mechanistic and therapeutic studies, particularly in dissecting epithelial–immune–stromal circuits relevant to CD. The identification of *Pcsk6+* enterocytes, given their known proinflammatory role in colitis, provides an entry point for studying epithelial drivers of immune polarization and mucosal injury.

## ACKNOWLEDGMENTS

We would like to thank the CWRU Genomics Core for sequencing the libraries generated during this study, and the Cleveland Digestive Disease Research Core Center (DDRCC)’s Histology/Imaging and Mouse Models Cores for histology/IHC services and provision of SAMP/AKR mouse strains, respectively. We would also like to acknowledge Daniele Corridoni and Greet De Baets for their helpful discussion of the human dataset. Many thanks to Katreya Lovrenert for uploading our data to GEO. Finally, we would like to thank the entirety of the Institute for Glial Sciences and the Cleveland DDRCC for productive discussion of this project.

## CONTRIBUTORSHIP STATEMENT

Conception and design: KL, BI, PM, MS, PT, FC

Data acquisition: KL, MS, BI, CDS, PM, BA, NK, AT, KC

Single-cell/nucleus data analysis: KL

GeomX analysis: BI, NK

Other data analysis and interpretation: KL, BI, CDS

Manuscript preparation: BI, KL

Critical revision of the manuscript: MS, FC, PT, TP

Study supervision: MS, FC, PT, TP

Funding: PT, FC, MS

## METHODS

### Animals

SAMP strains were described previously.[3] AKR/J mice (Strain #: 000648) were obtained from Jackson Laboratory and aged to 20 weeks in the Case Western Reserve University Digestive Diseases Research Core Center (CWRU DDRCC). All SAMP mice were bred and maintained on a mixed AKR/J genetic background in the animal facilities of the (CWRU DDRCC) under specific pathogen-free conditions, cohoused with AKR animals. Mice are routinely tested for pathogens. Experiments were performed in accordance with all current ARC and IACUC approved. All mice were observed for morbidity and euthanized when needed according to animal welfare. Cohorts included both male and female mice to promote sex-inclusive practice; no sex-based hypotheses or stratified analyses were prespecified. In total, 7 animals were used for snRNAseq, 13 animals for immunofluorescence and immunohistochemistry, and 8 animals for western blot. No animals were excluded from experiments. Since control (AKR) and experimental (SAMP) groups were defined by mouse strain rather than treatment, no randomization was used to allocate mice to groups.

### CitraPrep nuclei isolation for adult mouse small intestine

Mouse ileal tissue was dissected from 10cm proximal to the cecum up to the cecum and stereomicroscopy confirmed pathogenesis. Tissue was flushed with ice cold PBS to remove luminal contents, and snap frozen in liquid nitrogen. Snap frozen tissues were stored at -80°C until nuclei isolation. Nuclei prep was performed via the CitraPrep technique, as described previously [6]. In brief, frozen tissue was enzymatically and mechanically dissociated with glass dounce homogenizers (Wheaton Science Products, 357542), followed by separation via an iodixanol gradient. Nuclei were collected, washed and counted using a hemocytometer. 0.5U/uL RNAsin and 0.5U/uL SUPERase RNase inhibitors were included in all reagents and all steps were performed on ice.

### Single nucleus RNA-sequencing

Single nuclei suspensions for each sample were loaded into a separate well of a Chromium 10X Genomics single cell 3’ library chip, aiming to recover 15,000 nuclei (10X Genomics: Chromium Next GEM Chip G Single Cell Kit 1000127). Libraries were prepared as per the manufacturer’s protocol but with a modified fragmentation time of 4 minutes (10X Genomics 3’ GEM Library and Gel Bead Kit v3.1 1000128). All libraries were sequenced with paired-end reads following 10X Genomics guidelines on an Illumina Novaseq X Plus at the Case Western Reserve School of Medicine Genomics Core, with a goal of 50,000 reads per nuclei.

### Processing single nucleus RNA-sequencing

Sequencing data were aligned and counted using CellRanger v7.2.0 followed by CellBender v0.2.2, optimizing the number of epochs, fpr, and learning rate for each individual dataset based off ELBO learning curves and UMI decay curves [39]. CellBender uses a maximum-likelihood interference algorithm to filter ambient RNAs and barcode-swapped reads from the CellRanger’s raw count matrix output. The filtered dataset was then processed to remove nuclei with <250 or >8000 genes (online supplemental figure 1C). Only nuclei with fewer than 5%mitochondrial gene contamination were retained, which was the large majority (online supplemental figure 1D). A total of 73,156 nuclei passed quality control and doublet filtering with a mean and median detection of 2264 and 2044 genes per nucleus and a mean and median detection of 5969 and 4232 UMIs per nucleus. We next used DoubletFinder to selectively remove heterotypic doublets [40–42]. First, we generated artificial doublets from existing snRNAseq data before pre-processing the merged artificial-real dataset. We used the PC distance matrix to find the cell’s proportion of artificial k nearest neighbors to then rank and threshold according to the expected number of doublets with pANN threshold set to 0.10 and artificial doublets at 0.25.

The 7 mouse datasets were normalized using SCTransform with variable regression set to mitochondrial reads using the Seurat 5.1 package[43–47], followed by principal component analysis prior to clustering. We mathematically defined the optimal principal components (PC) to use first by calculating the lowest PC exhibiting >90% cumulative and <5% PC-specific variation. Then we found the highest PC whose percentage variation is >0.1% higher than the subsequent PC. The smaller of these two values was used as the optimal PC value for downstream analysis. Harmony [48] was used to batch correct by original sample identity. Unsupervised clustering resolution was initially set at 1 and adjusted until all clusters represented transcriptionally unique groups as determined by visualizing a heatmap of expression of the top 10 marker genes per cluster. Clusters with >50% of the top 50 DEGs associated with mitochondrial or ribosomal RNA (indicating RNA degradation) were removed, and normalization, batch correction, and clustering were repeated after their removal. Clusters were visualized by UMAP and clusters were annotated by cross referencing the top 100 markers per cluster with available literature, as well as by visualizing expression of key known marker genes of different cell types.

### Cell Proportion Analysis

Cell proportions were analyzed using the propeller method of the publicly available speckle package[49], which calculates sample-level cell type proportions, performs arcsin square root transformation, and then tests whether transformed proportions are significantly different in SAMP vs AKR using t-tests moderated using an empirical Bayes framework. Finally, false discovery rates (FDR) are calculated to account for testing across multiple cell types. Proportion changes with q<0.05 are considered statistically significant.

### Differential gene expression analysis

Gene expression differences were found with the DESeq2 pseudobulk method in Seurat, aggregating gene expression by cell type and by sample then comparing SAMP vs AKR. This method takes advantage of our high-powered dataset to identify consistent changes across biological replicates. Genes were deemed differentially expressed if they had an adjusted p-value < 0.05 and |Log_2_(Fold Change)| > 1. Functional enrichment analysis was performed using Metascape[50], which combines over 40 independent knowledgebases of functional enrichment, interactome analysis, and gene annotation in a single portal.

### Human scRNAseq analysis

Seurat v5.1 objects were created from publicly available processed data (matrix, features, and barcodes files) and metadata from Kong et al. on the Broad Institute’s Single Cell Portal (SCP1884).[10] Expression dot plots and module scores were calculated and visualized using built-in Seurat functions.

### CellChat Interactome Analysis

Cell-cell communication networks were inferred using the CellChat v2 R package.[28] CellChat predicts active signaling by integrating expression of ligands, receptors (including multi-subunit complexes) and cofactors of signaling pathways in their extensive database. Communication probability was estimated using a simplified mass-action-based model, employing the trimean method for calculating average expression levels per cell type. Global cell-cell communication networks were first calculated separately for AKR and SAMP cells, and then CellChat’s built-in differential signaling analysis pipeline was used to compare signaling pathways and interaction strength between the two groups.

### Fresh Fixed Paraffin-Embedding (FFPE)

Experimental mice were sacrificed and the terminal ileum was removed (10cm total length), opened longitudinally, rinsed in PBS, and submerged in 10% buffered formalin overnight. Formalin-fixed tissues were then washed, transferred to 70% ethanol, processed, and paraffin-embedded. Paraffin-embedded tissues were sectioned at 3–4 µm thickness, mounted on Superfrost Plus glass slides (Thermo Scientific, Logan, UT), and deparaffinized before immunostaining.

### Frozen tissue sectioning

Mouse ileal tissue was dissected from 10cm proximal to the cecum up to the cecum. Tissue was flushed with ice cold PBS to remove luminal contents, then cut open to a flat sheet and rolled into a Swiss roll before being fixed in 4% paraformaldehyde (PFA) overnight at 4°C. Fixed tissues were washed in PBS and moved to 30% sucrose in PBS and stored at 4°C overnight (until the tissue sank). Tissues were then embedded in Optimal Cutting Temperature Compound (OCT; Sakura, 25608-930), sectioned on a Leica CM1950 Cryostat at 15-20 um onto slides, and stored at -80°C.

### Histological evaluation of inflammation

Paraffin-embedded mouse ileal sections were sectioned and stained with haematoxylin and eosin (H&E). H&E-stained ileal sections were evaluated in a blinded, semiquantitative manner to assess inflammation development across different layers of the intestine. The evaluation was based on the following scale (total: 0-13). Lamina propria: Inflammation scale of 0-4 and expansion scale of 0-1 across the 6cm of ileum. Submucosa: Inflammation scale of 0-4 and expansion scale of 0-1 across the 6cm of ileum. Muscularis propria: Inflammation scale of 0-2 and expansion scale of 0-1 across the 6cm of ileum.

### Immunohistochemistry

Immunohistochemical (IHC) staining was performed using unconjugated polyclonal anti-mouse antibodies for DCLK1 (1:100, ab31704; Abcam, Cambridge, UK) or ST2 (AF1004; R&D Systems). Deparaffinized tissue slides were incubated in normal serum for nonspecific blocking. Endogenous peroxidase activity was quenched using 1.75% H₂O₂. Following overnight incubation with primary antibody at 4°C, slides were incubated with a biotinylated secondary antibody (Vector Labs) and developed using the Vectastain ABC Kit (Vector Labs). Immunoreactive cells were visualized using a diaminobenzidine (DAB) substrate (Vector Labs), counterstained with hematoxylin, and mounted using 80% glycerol. All incubations were performed at room temperature unless otherwise specified. Negative controls were prepared under identical conditions without the primary antibody. Tuft cells were quantified as the average number of intact, nucleated DCLK1-positive cells across the entire collected ileal tissue, normalized per high-power field (20X). Each data point in the graph represents one mouse, with 3 AKR and 4 SAMP mice shown in total.

### Immunofluorescence

For immunofluorescent staining, either frozen (epithelial stain) or FFPE (stromal stain) slides were incubated overnight at 4°C in primary antibodies diluted in blocking solution: PBS+0.1% Triton X-100 (VWR) with 5% donkey serum (Jackson ImmunoResearch). Primary antibodies included: PCSK6 (anti-goat AF488, Novus Biologicals NBP1-05030), c-MAF (anti-rabbit AF647, Proteintech 55013-1-AP), IGF1 (anti-goat AF488, Biotechne AF791), COL1A1 (anti-rabbit AF594, Boster PA2140-2), ACTA2 (anti-mouse AF647, Abcam ab7817). Slides were washed and incubated with Donkey secondary antibody at 1:500 for 2 hours at room temperature. All secondary antibodies were purchased from Jackson ImmunoResearch. After secondary, slides were washed and nuclei were stained with DAPI (Invitrogen) and mounted in Fluoromount G (SouthernBiotech, epithelial stain), or stained and mounted with Vectashield HardSet Antifade Mounting Medium with DAPI (H-1500-10, stromal stain). Negative controls were prepared under identical conditions without the primary antibody. Slides from 1 SAMP and 1 AKR animal were imaged per stain.

### Epithelial crypt-villus unit isolation

SAMP (n=4) and AKR (n=4) mice sacrificed and 10 cm of the ileum, starting from the cecum, were dissected. Adipose tissue, Peyer’s patches, and stool flushed with ice-cold Ca/Mg-free PBS. Incubate tissue in Ca/Mg-free PBS + EDTA 2mM for 15 min at room temperature on a slow rocker with gentle inversions. Release fractions (F) with brief 10–15 s shakes: collect F1 (villus-rich), then after 5-min re-incubations collect F2 and F3, which are crypt–villus unit (CVU)-enriched. Immediately stop each fraction with an equal volume of PBS with Ca/Mg on ice and enrich CVUs by gravity settling 1–2 min (or ≤50 g for 60–90 s); remove the supernatant and gently resuspend pellets using wide-bore tips. Avoid strainers or vigorous pipetting to prevent shearing. Pool F2+F3 for downstream western blotting use.[26]

### Immunoblotting

Tissue lysates were processed and run through sodium dodecyl sulfate-polyacrylamide gel electrophoresis (SDS-PAGE). Briefly, lysates were boiled with SDS-PAGE buffer for 5 minutes. Horseradish peroxidase-conjugated β-actin antibody was from Santa Cruz Biotechnology (Cat# sc-47778) and used in 1:5000 dilution (Cleveland, OH. PCSK6 antibody from Novus Biologicals (Cat# NBP1-05030) and used in 1:500 dilution (Cleveland, OH). Horseradish peroxidase-conjugated anti-goat IgG from R&D Systems (Cat# HAF017) was used in 1:5000 dilution (Cleveland, OH). The membrane was probed with β-actin as loading control and an image is shown.

### GeoMx whole transcriptome atlas

Four μm thick formalin-fixed paraffin embedded (FFPE) tissue sections were deparaffinized and washed prior to antigen retrieval in Tris-EDTA (pH 9.0) (eBioscience). RNA targets were then exposed using proteinase K (Thermo Fisher) and postfixed in 10% neutral buffered formalin. Slides were then incubated with GeoMx® whole transcriptome atlas probes at 370C overnight in a hybridization chamber (Boekel Scientific). Stringent washes in SSC (Sigma-Aldrich) and 100% deionized formamide (VWR) were performed to remove off-target probes. The slides were then stained with fluorescently labeled morphology markers: PanCK (AF488; 1:400; Novus), Syto83 nuclear stain (1:25; Invitrogen), Anti-alpha smooth muscle actin (AF594; 1:200; Abcam), and CD45 (AF647; 1:100; Cell Signaling Technologies). Following the manufacturer’s instructions, the slides are then loaded into the GeoMx® platform for imaging and barcode acquisition. Small intestinal AKR (control), SAMP non cobblestone (NC), and SAMP cobblestone (C) areas were identified and marked as regions of interest (ROI) for barcode acquisition. Library preparation was performed using a manufacturer-supplied PCR mastermix and well-specified indices to index and amplify the collected regions of interest. Barcodes were then pooled and purified using AMPure XP beads and ethanol washes. A Bioanalyzer DNA high sensitivity trace was used to assess library quality. Samples were sequenced on an Illumina platform. Quality control, normalization, and statistical tests were conducted on datasets for each region of interest.

### Statistical analysis

Statistical significance was defined as p-value or q-value < 0.05. This value was determined by moderated t-test (proportion analysis), Wald test (DESeq2), two-sample t-test (IHC/IF/disease scoring), or ANOVA with Tukey HSD (human module scores), with FDR discovery employed for all tests of sequencing data with multiple comparisons. In CellChat, significant interactions were identified by a statistical test that randomly permutes cell type labels. Sample sizes for all experiments were determined by consideration of effect (cell # or WB intensity) size and variance, and the assumptions of each test were met by the data.

## Code availability

Data analysis was performed with publicly available packages by following available tutorials and vignettes.

## Data Availability Statement: Data are available in a public, open access repository

All datasets generated from this work are included as Supplemental Tables in the Supplementary Materials. Fastq files and processed sequencing datasets have been submitted to the Gene Expression Omnibus (GEO) and will be uploaded pending NCBI approval (delayed during the U.S. government shutdown). Since we cannot yet provide an accession number, we suggest searching author names on the GEO portal or reaching out directly to corresponding authors.

## SUPPLEMENTARY FIGURES

**Figure S1:**
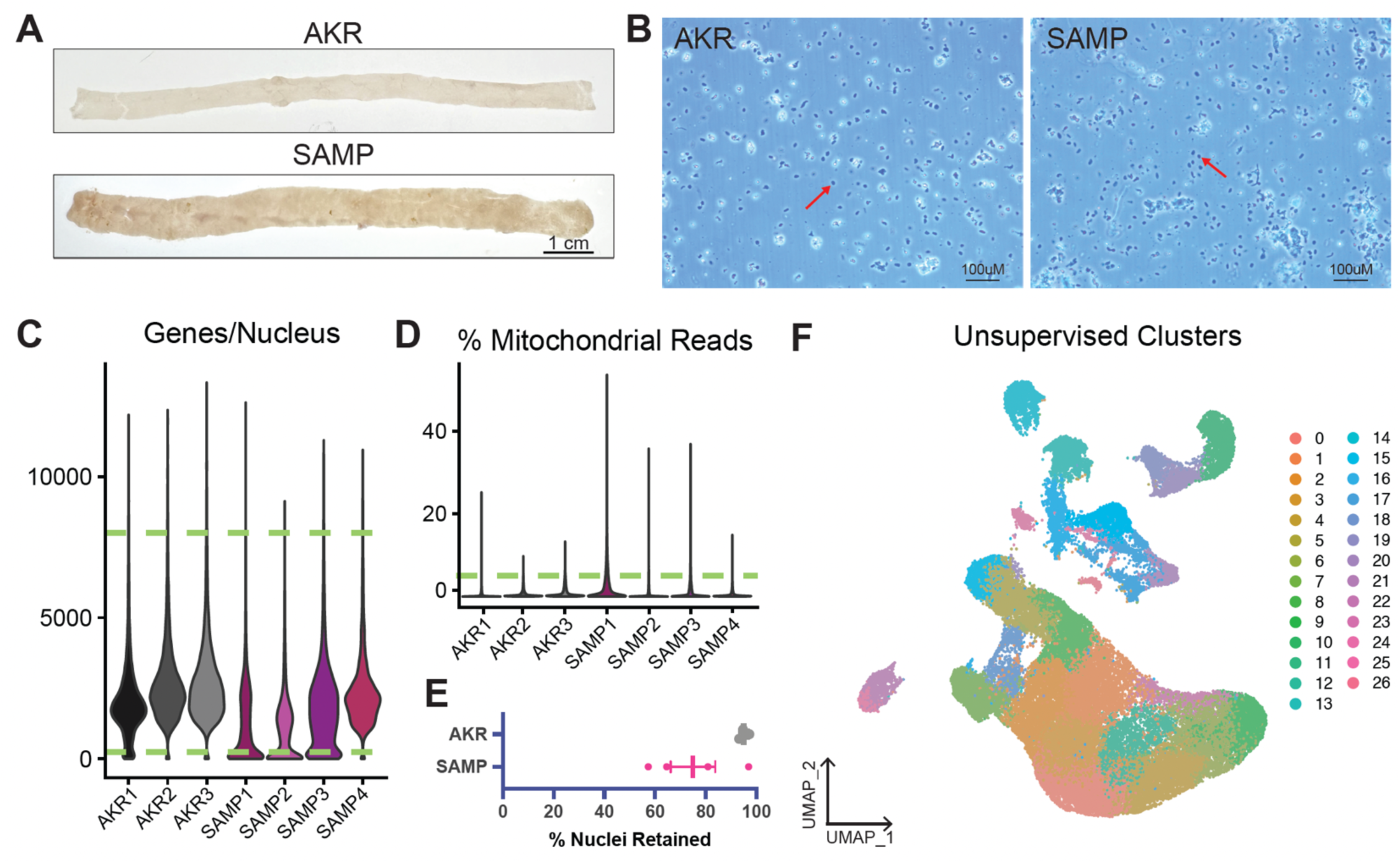
Quality control, sample integration, and unsupervised clustering for the whole-ileum single nucleus atlas. (A) Gross images of 20-week-old AKR and SAMP ileal at dissection. Scale bar represents 1cm in length. (B) Brightfield images demonstrating preserved nuclear integrity post-isolation in AKR (left) and SAMP (right); red arrows highlight representative nuclei. (C) Violin plots showing the distribution of genes per nucleus for each sample after CellBender-assisted preprocessing; nuclei with >8,000 or <250 genes per nucleus (dotted lines) were excluded. (D) Violin plots showing the distribution of mitochondrial read percentage per nucleus by sample; nuclei with <5% mitochondrial reads (dotted line) were retained (vast majority). (E) Nuclei retention rates after QC filters. (F) Unsupervised clustering of the 58,349 QC-retained nuclei projected in UMAP space.

**Figure S2.**
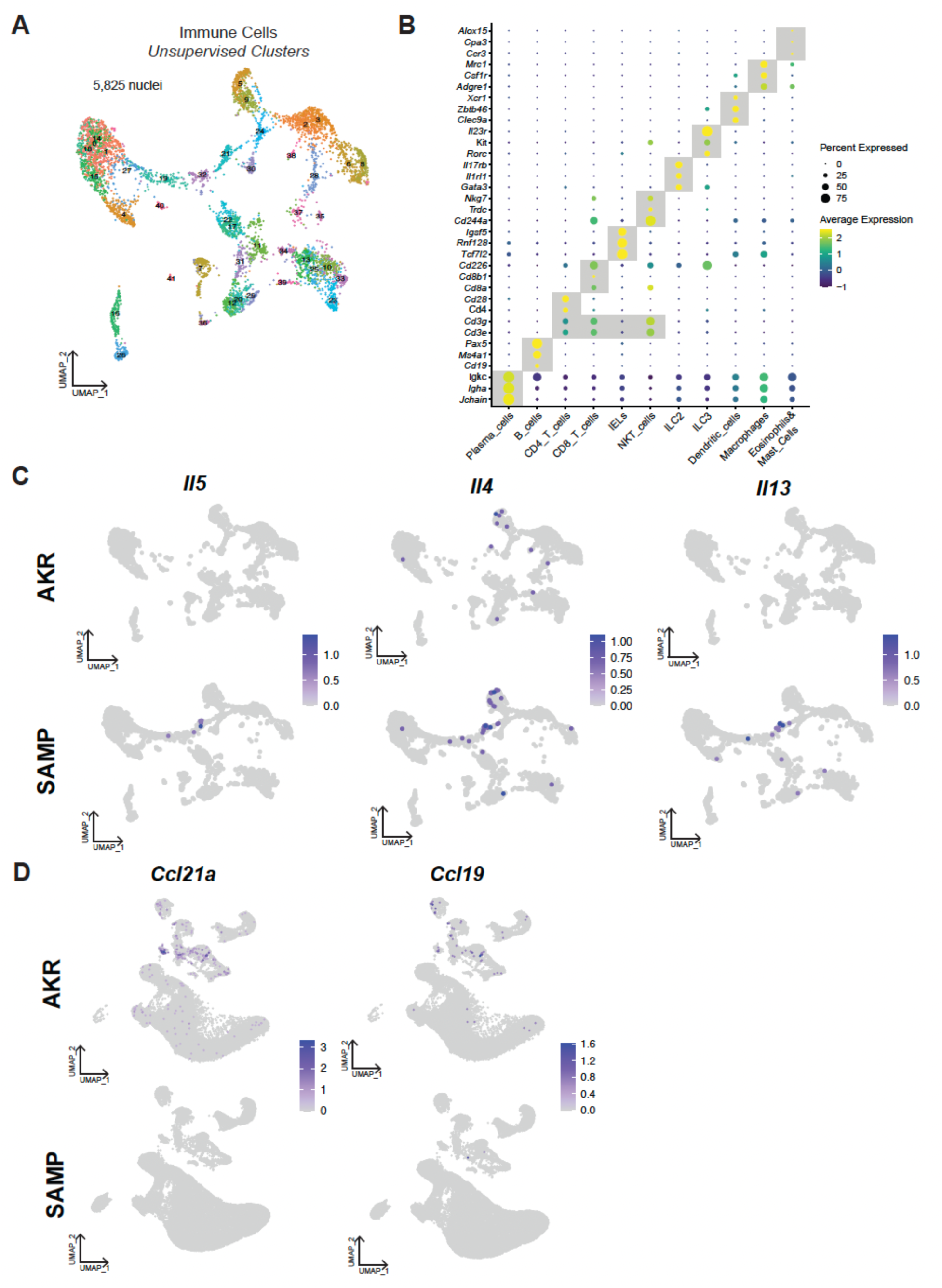
Immune compartment—marker validation and cytokine polarization. (A) Immune cells in UMAP space, colored by unsupervised clusters (5,825 nuclei). (B) Dot plot of cluster-defining markers confirming lineage identities across immune populations. (C) Strain-stratified feature plots for the canonical type-2 cytokines, *Il5*, *Il4*, and *Il13*. (D) Feature plots showing normalized expression of *Ccl21a* and *Ccl19* across the integrated atlas in UMAP space (color scale denotes expression level).

**Figure S3:**
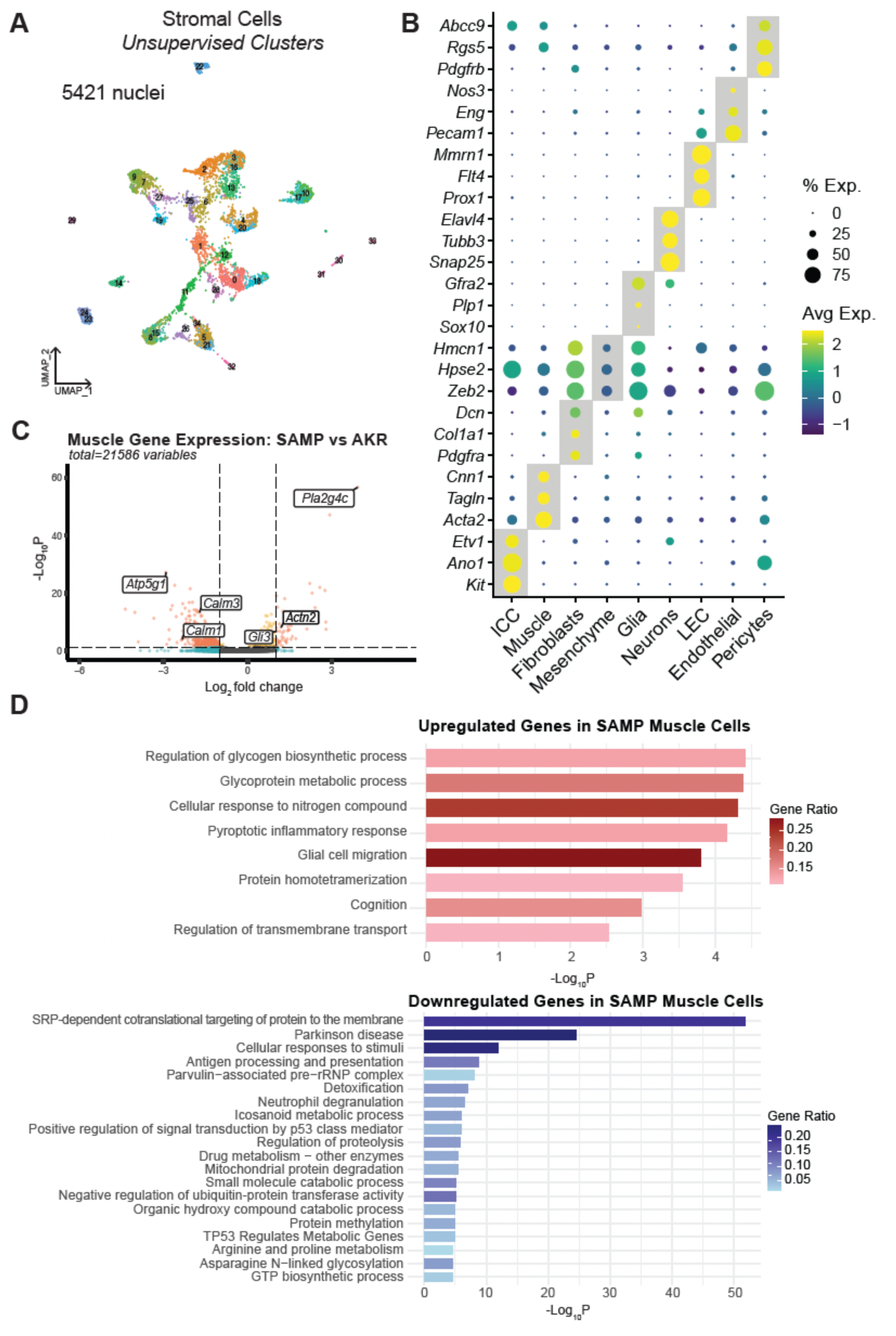
Stromal compartment—marker validation and muscle-cell transcriptional programs. (A) Unsupervised clustering of all 5,421 stromal cell nuclei from all samples, visualized in UMAP space. (B) Dot plot of cluster-defining markers confirming stromal lineage identities. (C) Differential gene expression in muscle cells of SAMP vs AKR mice. (D) Functional enrichment of upregulated and downregulated genes in SAMP muscle cells using Metascape.

**Figure S4:**
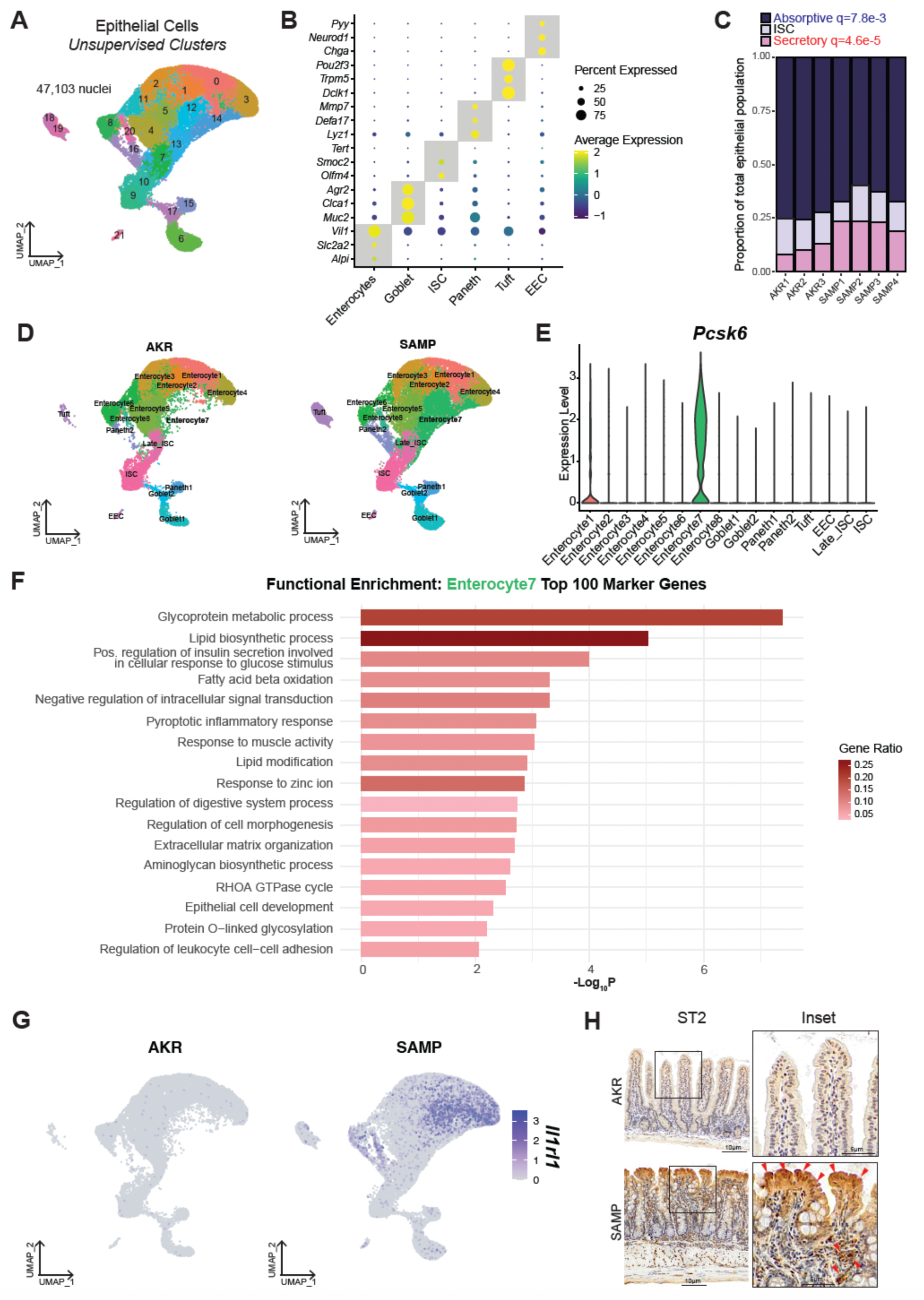
Epithelial compartment—composition, subcluster annotation, ST2/Il1rl1 features, and Enterocyte7 pathway enrichment. (A) Epithelial-only UMAP colored by unsupervised clusters (total nuclei indicated; e.g., 47,103 after QC). (B) Dot plot of canonical epithelial markers validating lineage identities. (C) Sample-level composition of the epithelial compartment across AKR1–3 and SAMP1–4, summarized as proportions of absorptive, secretory, and ISC categories. (D) Epithelial UMAPs annotated with subclusters for AKR and SAMP (Enterocyte1–8, Goblet1–2, Paneth1–2, tuft, EEC, ISC/late-ISC. (E) Distribution of *Pcsk6* expression across epithelial subclusters. (F) Functional enrichment of the top 100 marker genes for the Enterocyte7 subcluster using Metascape. (G) *Il1rl1* expression in AKR and SAMP epithelium plotted separately for visualization of this differentially expressed gene (DEG) in UMAP space. Color represents normalized expression compared with all genes across all epithelial cells. (H) Immunohistochemistry for ST2 (encoded by *Il1rl1*) in AKR and SAMP ileum with inset magnifications; scale bars as labeled.

**Figure S5.**
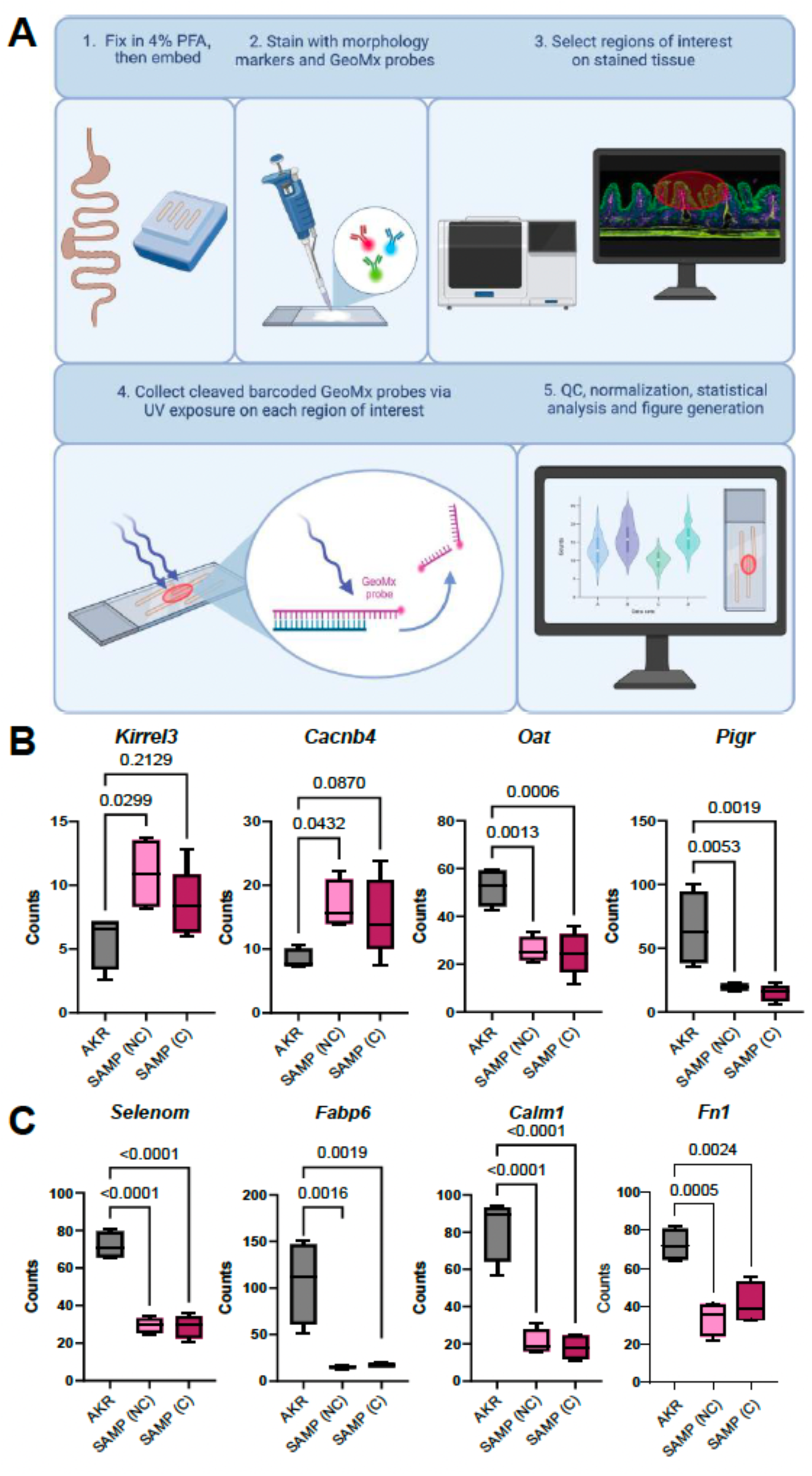
Mouse spatial transcriptomics— compartment-specific counts across epithelial (PanCK+), immune (CD45+), and smooth muscle (SMA+) regions. (A) Overview of spatial transcriptomic pipeline from tissue collection through analysis. (B) Gene expression counts from CD45+ immune regions of interest, displayed with group medians/interquartile ranges; exact P values reported on panel. (C) Gene expression counts from SMA+ stromal compartment regions of interest, displayed with group medians/interquartile ranges; exact P values reported on panel.

